# Spatial organization of assemblies of protein complexes by colocalization analysis

**DOI:** 10.1101/2025.10.27.684783

**Authors:** Daniel H. Orozco-Borunda, Antonio Martinez-Sanchez, Vladan Lucic

**Author notes:** Corresponding author: Vladan Lucic.

## Abstract

The precise localization of multiple species of proteins and protein complexes organized in nanodomains and large, non-periodic protein assemblies has been shown to be important for cellular function. Cryo-electron tomography is uniquely suited to visualize such assemblies in situ, with single nanometer precision, together with their cellular environment. Here we present in detail and characterize a parameter-free, second order point pattern analysis method that is applicable to cellular cryo-electron tomograms, and provides quantitative and statistical characterization of the colocalization between two or more distinct groups of complexes, such as those that comprise protein assemblies. By numerical and analytical calculations, we show that this colocalization method can correctly detect and distinguish different forms of point pattern interactions between complexes, and therefore identify specific types of spatial distribution of complexes, which result from multiple biochemical interactions and diffusion in cellular environments. We also present image processing tasks that precede the colocalization to facilitate its applications to cellular junctions and other biological systems.

## 1 Introduction

Most proteins are not randomly dispersed within cells, but are located in distinct cellular compartments, forming macromolecular complexes comprising core molecular components, as well as transiently bound subunits that modulate their function. Furthermore, proteins and protein complexes are also organized in a diverse group of nanodomains, loosely organized and dynamic protein assemblies comprising multiple complexes, such as lipid rafts (Ma et al., 2015; Levental et al., 2020), and Ca^2+^ nanodomains at the presynaptic terminal (Wang and Augustine, 2015). Synaptic nanodomains, which contain proteins residing in pre- and postsynaptic neurons and may colocalize across the synaptic cleft, were first detected by super-resolution fluorescence microscopy and subsequently imaged by cryo-electron tomography (cryo-ET) (Perez de Arce et al., 2015; Tang et al., 2016; MacGillavry et al., 2013; Nair et al., 2013; Chamma et al., 2016; Maschi and Klyachko, 2017; Hruska et al., 2018; Glebov et al., 2017; Liu et al., 2020; Martinez-Sanchez et al., 2021). These show that assemblies comprising multiple species of protein complexes, which often emerge from colocalization of nanodomains, form functional modules and are critical in synaptic transmission and possibly other cellular processes (Biederer et al., 2017; Chen et al., 2018; Choquet, 2018; Scheefhals and MacGillavry, 2018; Zuber and Lucic, 2022).

The development of cryo-ET, which allows single nanometer resolution imaging in three dimensions (3D) of fully-hydrated, vitrified biological samples (Dubochet et al., 1988; Lucic et al., 2005; Beck and Baumeister, 2016; Oikonomou and Jensen, 2017), resulted in a considerable number of high resolution structures of proteins and protein complexes in situ (Turk and Baumeister, 2020; Ng and Gan, 2020). Furthermore, cryo-ET is uniquely suited to image entire cellular regions and provide a comprehensive and accurate representation of a large number of molecularly heterogeneous complexes integrated in their native environment. This offers possibilities to characterize the spatial organization of non-periodic large protein assemblies comprising a multitude of complexes within crowded cellular environments, in order to determine their contribution to cellular function.

Spatial point pattern analysis aims to characterize point distributions and provide statistical methods for distinguishing them from completely random distributions (Ripley, 1981; Wiegand and Moloney, 2004). Of particular importance in the cellular context are bivariate methods for detection of point pattern interactions because they indicate the presence of direct or indirect biochemical interactions or other cellular mechanisms that organize protein complexes. To clarify the terms coming from different fields, proteins and protein complexes (here together termed complexes) are mathematically represented by spatial points, while distinct groups of complexes (for example distinguished by their molecular identity) form the corresponding point patterns, also called particle sets in cryo-ET.

Here, we present a detailed characterization of the colocalization method. It is used to analyze the distribution of pair distances between particles of two or more sets and detects point pattern interactions between them, thus belonging to the second order, bi- (or multi-) variate point pattern methods (Wiegand and Moloney, 2004; Martinez-Sanchez et al., 2022). We derived analytical solutions, as well as numerically calculated colocalizations for several well defined, synthetic point patterns and determined their statistical significance, in order to properly characterize the colocalization method and determine rules for interpreting the results. This method was previously applied to experimental cryo-ET data to detect trans-synaptic assemblies at neuronal synapses (Martinez-Sanchez et al., 2021). In addition, we describe and give usage tips for all other tasks comprising the colocalization workflow, which precede the colocalization task and ensure that the colocalization is set up properly (Figure 1).

**Figure 1.**
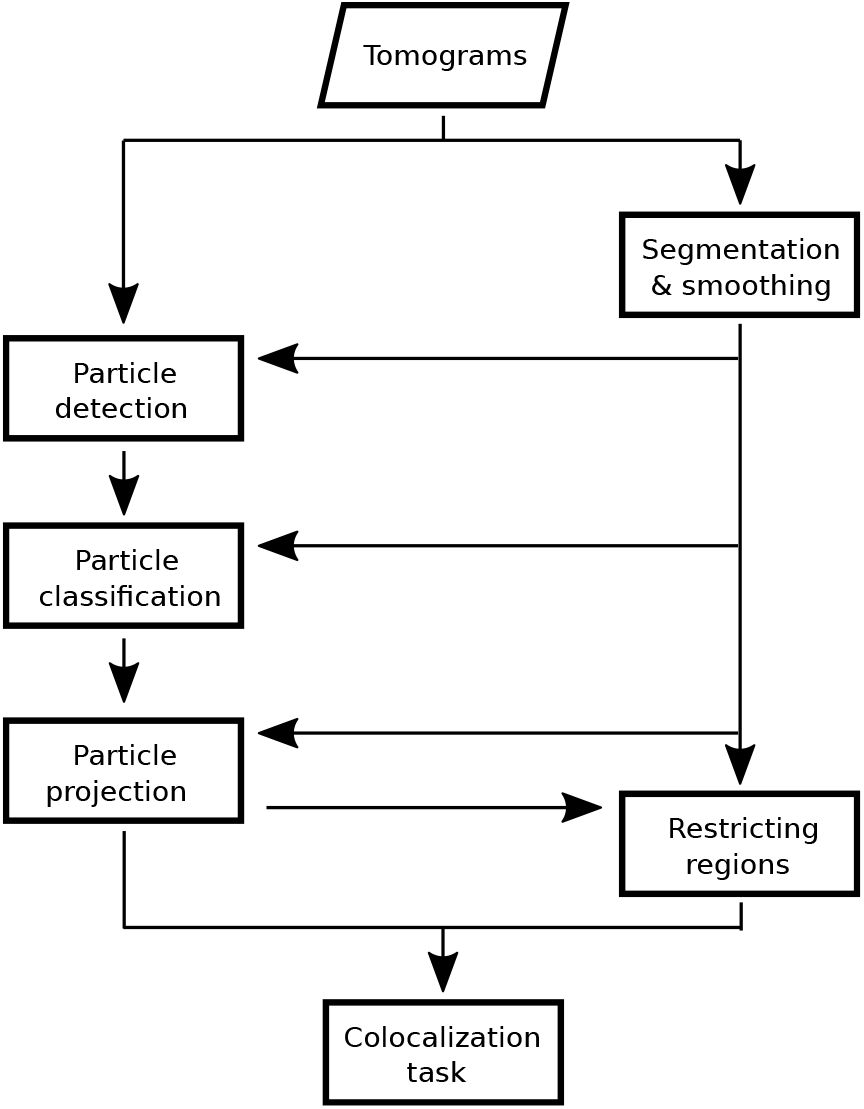
Colocalization workflow scheme. Tasks comprising the workflow are indicated by rectangles.

## 2 Results

### 2.1 Colocalization task definition and theoretical considerations

#### 2.1.1 Colocalization task definition

The colocalization task aims to elucidate the joined organization of two or more experimentally determined particle sets and especially to detect and quantify interaction between particle sets, by analyzing their spatial location ((Martinez-Sanchez et al., 2021)). Specifically, given the precise location of two particles sets here called *P* and *Q* (particles can be proteins or protein complexes), and a colocalization distance *d*, the number of colocalizations between sets *P* and *Q* is defined as the number of particles in *P* that have at least one particle of *Q* located at the Euclidean distance smaller or equal to *d*. Clearly, colocalization defined in this way is not invariant under the exchange of the order of the sets.

The Monte Carlo method is used to calculate the statistical significance of the colocalization, that is to determine whether the number of colocalizations between given particle sets is significantly higher than the null model. We distinguish between”experimental”particle sets whose colocalization is examined (here called *P* and *Q*), and the simulated, completely randomized particle sets (simulated *P* and simulated *Q*) that correspond to *P* and *Q* and are generated during Monte Carlo tests.

Two null models were designed in order to decouple clustering (univariate heterogeneity) from the interaction between particle sets (bivariate heterogeneity) (Wiegand and Moloney, 2004), resulting in two Monte Carlo tests. Namely, in the”standard”null model particles of the first experimental set are kept fixed and randomized particle sets corresponding to the other particle set are used (experimental *P* and simulated *Q*), while in the “alternative”null model the first model is randomized and the second is fixed (simulated *P* and experimental *Q*) ((Wiegand and Moloney, 2004; Martinez-Sanchez et al., 2021)), The two p-values can be averaged (weighted by the number of simulations) to obtain the combined p-value. In both cases, multiple simulated sets are generated in order to obtain the probability that the null model is correct, resulting in the standard and alternative p-values.

The above defines the colocalization between two particle sets (2-colocalization). The 3-colocalizations (and higher) are defined by calculating the number of particles of the first set that have at least one particle from each of the other sets within its neighborhood *d*. For the standard null model the first particle set is fixed and all other are randomized, while for the alternative null model the first is randomized and all other are fixed. Consequently, the order of particle sets matters in that it is important which set is the first, but the ordering of the other sets is irrelevant (colocalization between sets *P*,*Q* and *R* is the same as *P*,*R* and *Q*, but different from *Q, P* and *R*).

Together, the colocalization task requires particle locations (coordinates) so a particle set is simply a point pattern. Furthermore, it requires tomograms showing regions (boundaries) where the (projected) particles are located. The same regions are then used to distribute the randomized particle sets, so that they fully correspond to their respective “real” particle sets and make statistical comparison possible. Particle sets can be present in one tomogram, or split in separate units (subsets) across multiple tomograms or different parts of tomograms. In any case, each particle set unit has to have an associated spatial region.

#### 2.1.2 Analytical solutions

Based on the above colocalization definition, we derived analytical solutions for the number of colocalizations between particle sets (Eq 4, see the Appendix).This equation is valid for the colocalization of two or more particle sets. It is a function of the colocalization distance and depends on the following parameters specified by the colocalization problem: number of particles of the sets, the area over which they are distributed and the intersection of all particle set areas.

To account for the effects caused by the boundaries of the particle set areas, we introduced two parameters whose values have to be determined heuristically: (i) the effective dimensionality *ξ*, which for values smaller than 2 compensates for the overestimate of the number of colocalization by the analytical formula caused by the particles of the first set that are located to the area border, and (ii) the overlap area factor *α*, which for values greater than 0 compensates for the underestimate by the formula caused by the fact that for a colocalization *P* -*Q*, when the area of *Q* is inside the area of *P*, particles of *P* located outside the intersection area also contribute to the colocalization (Suppl. figure S1A).

Finally, we introduced a heuristic parameter *β*, which for values greater than 1 mimics the interaction between particles of different sets.

### 2.2 Numerical solutions

Here we numerically calculate the number of colocalizations between different particle patterns (distributions) for a range of colocalization distances, determine their statistical significance, and compare them with the analytical results. These patterns include completely random particle distributions, univariate distributions, as well as interacting (bivariate) patterns where particle locations of different sets are correlated with each other. The latter comprise patterns arising from global interactions where two particle sets are clustered over similar regions, or from local interactions where individual particles of one set are localized in the vicinity of the other set, as explained below.

Graphs presented here shoe the standard and the alternative p-values. Because the combined p-value can be simply inferred as the average between the standard and the alternative p-values, the combined p-values are not shown explicitly. The number of particles in a pattern, the area of the regions where they were distributed, the pixel size, the range of colocalization distances, and the particle exclusion distance were chosen according to the previously obtained cryo-ET results ((Martinez-Sanchez et al., 2021; Papantoniou et al., 2023)).

#### 2.2.1 Completely random patterns

As expected, colocalization between two completely random particle patterns was not significant (Figure 2A). While the analytical solution (without corrections) (Eq 1) showed a good agreement with the numerical result, correcting the effective dimensionality (*ξ* = 1.96) (Eq 4) provided a noticeably better fit.

**Figure 2.**
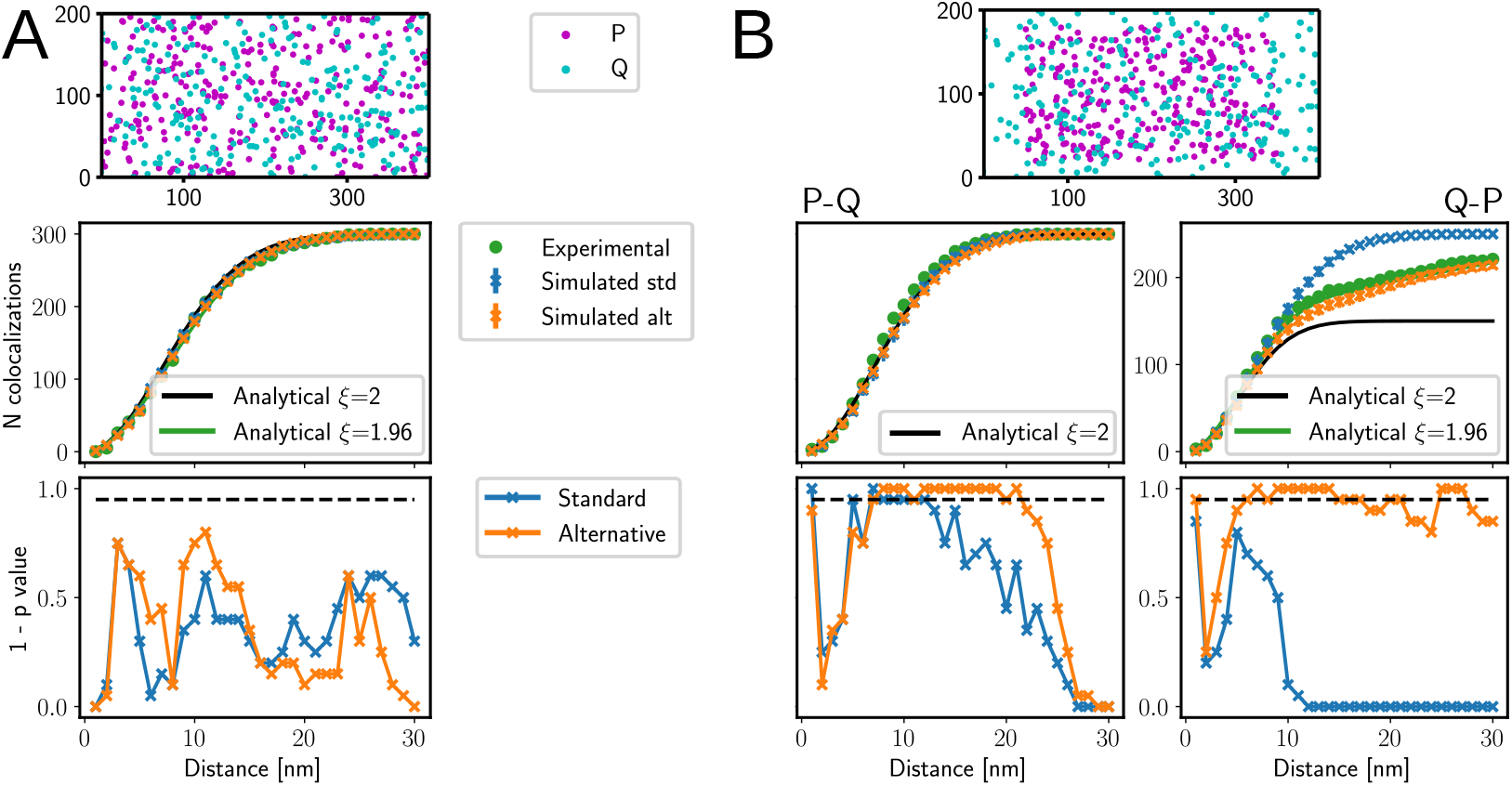
Colocalization between non-interacting particle sets *P* and *Q*. (A) Completely random, independent sets. (B) Colocalization between a set clustered on 60% of the entire area and a completely random set. (A-B) Particle distribution is shown on top, number of colocalizations in the middle and p-values at the bottom. Horizontal dashed line shows 0.05 confidence limit. The colocalization order is specified above graphs. Legends shown between panels apply to both.

Next, we changed the shape of the rectangle where the particles were distributed to make it extremely elongated, from 200×400 pixels to 20×4000 pixels, while keeping the area and the number of particles the same. The numerical result showed a decreased number of colocalizations that did not agree with the analytical solution (Eq 1). However, further decreasing the effective dimensionality parameter (*ξ* = 1.85) provided a better agreement (Fig. S2A).

Together, these observations showed that the number of colocalizations between random patterns was not significant. Also, adjusting the effective dimensionality parameter *ξ* was needed for the analytical solution to provide a proper boundary correction.

#### 2.2.2 Univariate patterns

We proceeded by generating clustered particle patterns, where the same number of particles used for patterns *P* and *Q* was randomly distributed within a smaller area (termed clustering region).

Colocalizations between particle set *Q* distributed over the entire area (*Q* full) and *P* distributed over 80%, 60% and 50% of the entire area (*P* clustered) showed that the number of experimental colocalizations was significantly lower than the number obtained from the standard simulations (Figure 2B, Suppl. figure S2C,D). This can be understood by observing that particles from set *Q* that are located outside the clustering region of set *P* do not contribute to the colocalization in the experimental case, may contribute to the normal simulations because the simulated set *P* spreads outside the entire area. Calculating the same colocalizations, but using particle set *R* instead of *Q*, which has 5 times smaller number of particles than set *Q*, supported the conclusions we reached for *P* and *Q* (Suppl. figure S2E).

Inverting the order of the particle patterns, that is calculating colocalization between *P* clustered and *Q* full, deviated only slightly from the null model and reached significance at few isolated distances (Figure 2B, Suppl. figure S2C-E). This was likely caused by the stochastic distribution of the experimental particle set because the expectation of the experimental and standard test colocalizations are the same.

The analytical solution (Eq 1) agreed with the numerical results for the colocalization between *P* clustered and *Q* full, However, colocalization between *Q* full and *P* clustered produced very different results. Namely, the experimental and the number of colocalizations from the alternative simulation far exceeded values obtained from the uncorrected analytical solution (Eq. 1) and the difference increased with the increased colocalization distance (Figure 2B). This indicates that the discrepancy was due to the boundary effect that can be compensated by an increase of the effective number of particles of *Q* that are within the area of *P*. Indeed, setting parameter *α* = 0.015 in Eq. 4 removed this difference, thus justifying the rational for introducing this parameter. Furthermore, standard simulation completely agreed with the analytical solution In addition, we generated another type of univariate patterns, where particles form a more regular (less clustered) pattern than the completely random one, by imposing a minimal exclusion distance on 5 nm between particles. The number of colocalizations obtained for the two cases when a minimal exclusion distance was imposed on one set and the other was completely random were very close to the analytical solution for completely random patterns and the combined test was not significant (Suppl. figure S2B).

Together, our results show that univariate clustering of the second set in colocalization, but not of the first set, can be detected by a significantly higher colocalization of standard simulations than of the experimental case. This argues that useful information can be obtained when both particle set orders are considered in colocalization. Furthermore, we showed that a small exclusion distance dos not lead to a significant colocalization. Also, adjusting the effective number of particles parameter *α* correctly accounted for boundary effects arising from particle clustering.

#### 2.2.3 Global interactions

Next we investigated colocalizations when particle sets are co-clustered, that is both sets are clustered over the same or similar areas. We call these globally interacting sets because they are not independent from each other as their clustering regions are related, even though the individual particles were randomly distributed within these regions.

Colocalizations between clustered particle sets *P, Q* and *R*, with completely overlapping clustering regions covering 60% or less of the entire area, showed significance for the alternative test over almost all colocalization distances, for both orders of particle sets (Figure 3A, Suppl. figure S3A, B). The standard and combined tests were significant for a large range of distances, typically from small distances up to those where the number of colocalizations reached saturation. The analytical solution with corrected effective dimensionality showed very good agreement with the numerical one (*ξ* = 1.96). Furthermore, the number of colocalizations obtained for the standard test was larger than for the alternative test for a range of colocalization distances. In addition, the experimental number of colocalizations was significantly lower than that obtained for standard simulations at distances where both colocalizations where in saturation (Figure 3A, Suppl. figure S3A-C), indicating that this effect might be a consequence of a stochastic particle distribution.

**Figure 3.**
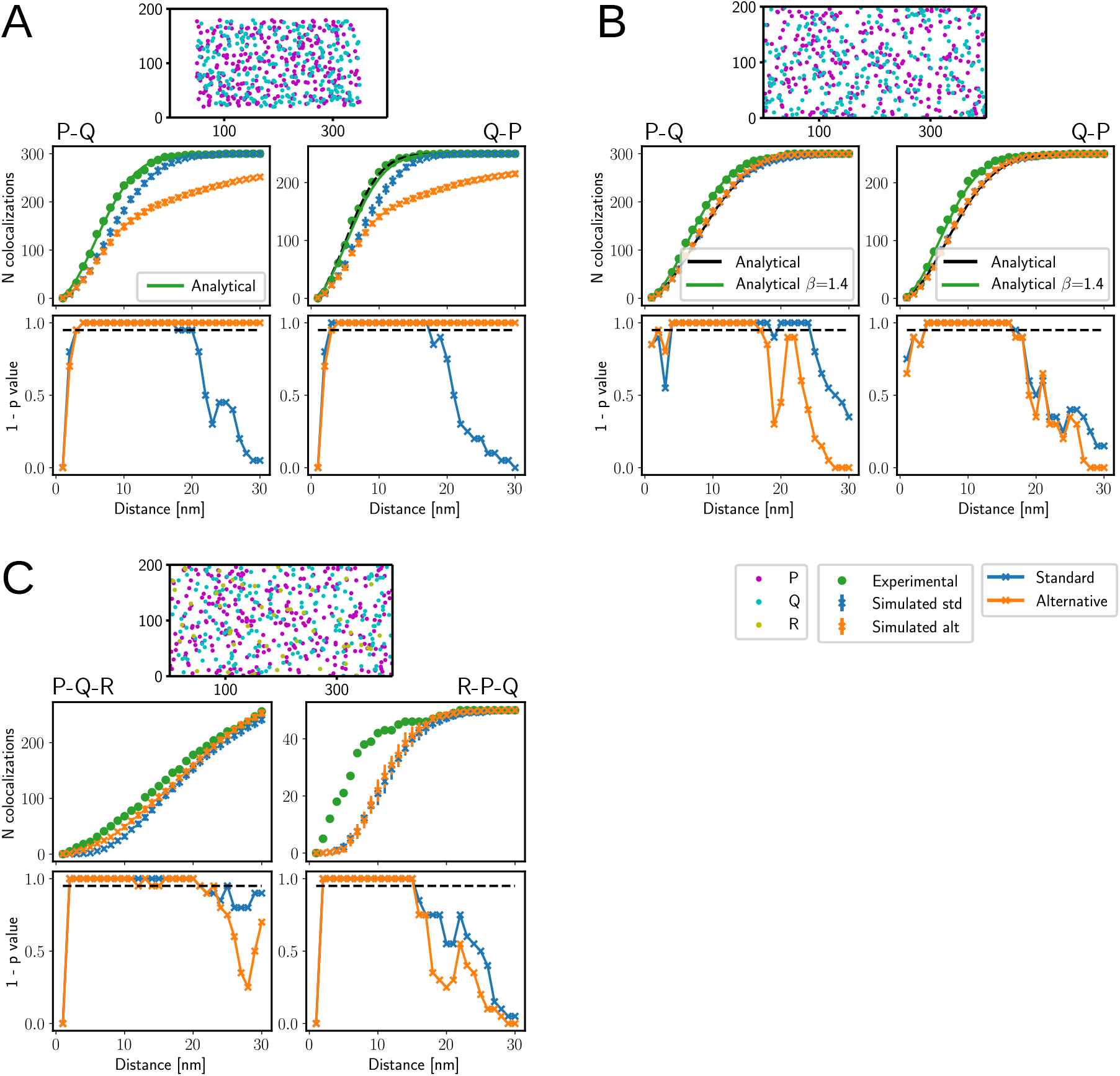
Colocalizations between interacting particle sets. (A) Globally interacting sets, both are clustered on completely overlapping regions covering 60% of the entire area. (B) Locally interacting sets, 1-to-1 mode, the interacting subset contained 30% particles of *P*, interaction distance 10 nm. (C) 3-Colocalization, local interaction, common interacting subset variant, the subset contains 10% particles of *P*, interaction distance 10 nm. (A-C) Particle distribution is shown on top, number of colocalizations in the middle and the p-values at the bottom. Horizontal dashed line shows 0.05 confidence limit. The colocalization order is specified above graphs. Legends shown between panels apply to all panels.

Separating clustering regions so that they overlap only partially resulted in a reduced range of distances at which colocalization was significant (Suppl. figure S3C). In addition, for larger colocalization distances, the number of experimental colocalizations was significantly smaller than that of standard simulations.

When clustering regions covered 80% of the entire area, the alternative test showed significance for a range of distances, but the standard test only for few distances (Suppl. figure S3D). Here also, standard simulations resulted in a higher number of colocalizations than the alternative simulations, for a majority of colocalization distances.

Therefore, even modest co-clustering resulted in robust, significant colocalizations, for both orders of particle sets and at distances where the number of colocalizations was not saturating. In all cases, except for small distances, standard simulations resulted in a larger number of colocalizations than alternative simulations.

#### 2.2.4 Local interactions

To generate locally interacting particle sets, we started from a random set, chose a subset of this set (interacting subset) and generated a dependent set in one of two ways. In the 1-to-1 mode, one particle of the dependent set was placed in the neighborhood of each particle of the interacting subset, thus making 1-to-1 interactions (Suppl. figure S1B left). In the multimeric mode, a subset of the dependent set was distributed over neighborhoods of all interacting subset particles (Suppl. figure S1B right). In the latter case, some interacting subset particles did not interact with any, while some interacted with multiple particles from the dependent set.

Colocalizations between clustered particle sets *P Q* and *R* was significant according to all three tests, regardless of the order of particle sets, roughly up to distances where colocalization reached saturation (Figure 3B). This was the case for the 1-to-1 mode for as low as 15% of the particles of the first set interacted with the second set, at interaction distances of 10 and 5 pixels (Suppl. figure S4A-C. In the multimeric mode, significance was reached when 50% or more particles were selected for interactions (Suppl. figure S4D). In all cases, contrary to the globally interacting case, standard and alternative simulations resulted in similar number of correlations.

In addition, we determined colocalization of particle sets that interacted both locally and globally. The results showed contributions from both interactions, as there were prominent differences between the experimental number of colocalizations and the standard simulations, as well as between the standard and the alternative simulation (Suppl. figure S5).

Analytical solution without the interacting factor fit both simulation results very well (Figure 3B). Adjusting the interaction parameter *β*, accounted well for the difference generated by the local interactions.

Together, we showed that a careful investigation of colocalization results allows detecting local interaction between two point patterns both in the presence or without global interaction (clustering).

#### 2.2.5 Colocalization between three patterns

To investigate particle interactions between three sets (3-colocalizations), we generated interacting particle sets in two ways. In the first variant (common interacting subset), the same interacting subset of *P* was used to generate (the interacting sub-) sets of *Q* and *R* (Suppl. figure S1C left). In the second variant (independent interacting subsets), two different interacting subsets of *P* were selected, one was used to generate set *Q* and the other one *R* (Suppl. figure S1C right). In both cases, particle set *P* was completely random, and sets *Q* and *R* were generated so that they locally interact with *P* in 1-to-1 mode, in the same way as for (2-)colocalizations described in the previous section.

In the common interacting subset variant, the colocalization showed strong confidence for both standard and alternative null models, over a range of colocalization distances, when even just 10% of particles from *P* formed the interacting subset (Figure 3C, Suppl. figure S6A). The independent interacting subsets variant showed a weaker correlation, reaching significance as low as 15% of particles from *P* in the interacting subset (Suppl. figure S6C, D). As expected, colocalization between three completely random and independent patterns *P, Q* and *R* did not reach significance for any of the tests (Suppl. figure S6B). In all cases, colocalization was calculated for two different particle set orders, *P* -*Q*-*R* and *R*-*P* -*Q*.

The analytical solution fit the completely random case as well as all simulations (Suppl. figure S6A). Even with the optimal choice of the interaction parameter, the analytical solution only qualitatively agreed with the colocalization results for interacting sets *P* -*Q*-*R*, while the colocalization *R*-*P* -*Q* showed a very good fit. This indicates that, depending on the number of particles, 3-colocalization interactions may or may not be modeled by a single interaction parameter.

In short, the colocalization procedure detected interactions between three particle sets even when 10% of the particles were interacting, and showed an increased sensitivity when the same subset of one particle set interacted with the other two sets.

#### 2.2.6 Guidelines for interpreting colocalization results

Based on the above results, we here provide guidelines for interpreting colocalization results.

In general, we suggest calculating colocalizations for different orders of particle sets because it allows distinguishing interactions between two sets from univariate distribution of one set. Also, significant results obtained for colocalization distances where the number of colocalizations is in saturation, should be interpreted with great care, or not considered at all, to avoid false positive results.

Particle interaction, both local and global, is recognized by a significant colocalization according to both null models (combined test) and in both orders of particle sets. The presence of global interaction (co-clustering), with or without local interaction is distinguished by a substantially higher number of colocalizations for standard than for alternative simulations.

A non-significant combined test result, even if one test showed significance, argues against particle interactions.

A significantly lower number of experimental colocalizations compared to standard simulations signifies that the second particle set is clustered. This can happen in two situations. First, if the same signature is observed for both particle set orders and it has already been determined that the particle sets interact, it is likely that the two particle sets are partially overlapping. That is, there is a component arising from global interaction (co-clustering), but also another one from clustering of one set that is independent of the other. Second, when this signature occurs for one, but not for the other order of particle sets, and particle interaction was not detected, it can be concluded that only one particle set is clustered (the particle set that was the second one according to the order that resulted in this pattern).

### 2.3 The colocalization workflow

The colocalization task requires precise locations (spatial coordinates) of multiple species of protein complexes, which are represented as point particle sets, as well as a segmented region where the particles are located. Here, we describe image processing tasks comprising the colocalization workflow that follow 3D reconstruction of tomograms and give usage tips, which, in our experience, ensure that particle sets and their locations are properly determined (Figure 1). Two of these tasks, detection of protein complexes and particle classification are only briefly summarized because they were presented before (Martinez-Sanchez et al., 2020). The tasks are presented in the order they are usually executed.

#### 2.3.1 Segmentation of membranes and subcellular regions

Segmentation of regions where protein complexes (particles) of interest are located is required for the colocalization task. These regions can be lipid membranes, intra- and extracellular regions, and other distinct, clearly visible cellular structures ((Volkmann, 2010; Martinez-Sanchez and Lučić, 2024)). Here we present usage hints from our experience with different tools to segment membranes suitable for for colocalization.

Manual segmentation, still a prevalent method in the field, is a demanding task that typically consists of tracing structures of interest on 2D tomographic slices. Amira™(Thermo Fisher Scientific)(Stalling et al., 2005; Kremer et al., 1996; Lamm et al., 2022) is a well-established GUI-based commercial software that contains many tools for precise pixel annotation and is often used in the field. The most prominent free, open source alternative in the 3D electron microscopy field, also GUI-based, is IMOD (Kremer et al., 1996; Danita et al., 2022). While 3D segments are represented by cubic pixels on a rectangular grid in Amira™, which allows the user to modify each pixel directly, IMOD segmentation is closer to vector graphics because it is based on geometrical objects like contours and surfaces (Suppl. figure S7A), In our experience, IMOD segmentation is less intuitive and has a steeper learning curve than the pixel-based segmentation, but once mastered may allow faster segmentations.

TomoSegMemTV software was specifically developed for automated membrane segmentation of cryo-ET images (Martinez-Sanchez et al., 2014). It is centered on Tensor voting algorithm, which uses robust second order derivative filter to detect membranes. Another automated tool that we used is Membrain-seg v2 (Lamm et al., 2024) is based on U-Net deep learning architecture (Shelhamer et al., 2017; Ronneberger et al., 2015). While it is possible to use the pretrained network to avoid time consuming training step, tomograms may need to be adjusted to the expected form.

Tools that explicitly use standard image processing methods (IMOD, TomoSegMemTV) have the advantage that one can understand the methods and optimize their usage. However these may be difficult to understand, so they are better suited for researchers having a mathematical or computational background.

Automated approaches provide good looking segmentation s relatively quickly (Fig S7A) (Lamm et al., 2022). However, a substantial subsequent manual segmentation is needed to correct automated segmentations to render them suitable for the colocalization workflow. Namely, segmented membranes need to be continuous, without holes, covering the entire regions where complexes of interest are present, and should not include other membranes or cellular structures.

The required segmentation accuracy depends on the further processing tasks. For example, while it is generally advisable to obtain an accurate segmentation, Morse theory based detection of membrane-bound complexes task (see below) allows membrane segmentation inaccuracy of the order of few nanometers because the 3D refinement step improves the localization accuracy of complexes. However, processing tasks that complement colocalization, such as the detection of membrane bridges and subtomogram averaging that includes a selective membrane filtering (Lucic et al., 2016; Orozco-Borunda et al., 2024), require a precise membrane segmentation.

In summary many existing segmentation tools can be used for the colocalization pipeline. The choice ultimately depends on the experience and the background of the researcher.

#### 2.3.2 Smoothing segmented regions by morphological filters

It is usually easier and less time consuming to segment regions starting from a tomogram that was binned by a factor such as 2 or 4 (e.g. at pixel size *>*1.5 nm). To use it for further processing, the segmentation has to be magnified to the original tomogram size, by replacing each pixel by a cube of size 2 or 4, respectively, which results in a serrated representation of surfaces that are not oriented along the major axes. Similarly, segmenting xy tomographic slices one by one often leads to the step-wise appearance in the z-axis direction (the staircase effect).

These effects can be mitigated by applying a series of binary morphological operations. For example, processing a segmentation that was binned by a factor of 2 using erosion(1), dilation(2) and erosion(1), or by a bin factor 4 using erosion(2), dilation(4) and erosion(2), where the numbers in parentheses indicate the number of iterations, can make segmented surfaces smoother (Suppl. figure S7B).

#### 2.3.3 Morse theory based detection of protein complexes - particle picking

The next task, particle detection, starts with a comprehensive template-free tracing of biological density by a discrete Morse theory based procedure ((Forman, 2002; Sousbie, 2011)) and a conversion of density traces to a spatially embedded graph representation ((Martinez-Sanchez et al., 2020)). The location of individual particles (molecular complexes) is detected by selecting subgraphs based on their location and distance to the previously segmented subcellular regions. For example, a subgraph representing a membrane-bound complex has to contain vertices belonging to both previously segmented membrane and a neighboring region (intra-, extracellular or lumenal). In addition, the direction of vectors normal to the region (typically a membrane) at locations in the vicinity of the particles is determined.

To obtain a more precise particle locations and membrane normals, particles are subjected to subtomogram 3D refinement where alignment angles are constrained and a high axial symmetry is imposed (such as C10). Because particles detected by the Morse theory method are highly heterogeneous in shape and size, the resulting average is expected to clearly show membrane and possibly a weak density along the normal direction. Observing a well resolved membrane is an important checkpoint in the processing pipeline. Finally, because the 3D refinement induces particle translations, a minimal inter-particle (exclusion) distance is imposed.

#### 2.3.4 Particle classification by Affinity propagation

Because the Morse theory particle detection is rather comprehensive, it generates a morphologically very heterogeneous set of particles, comprising complexes of many molecular species. Consequently, these particles need to be separated into classes that contain structurally similar particles and are sufficiently homogeneous to be further processed by the standard subtomogram 3D classification methods. This is a demanding classification requirement that may need different approaches for different complexes.

In our hands, the optimal method consisted of two steps, where particles are first rotationally averaged around their membrane normals, and then these axial averages are subjected to unsupervised clustering by Affinity propagation (AP) ((Frey and Dueck, 2007; Martinez-Sanchez et al., 2020)). We note that this method is suitable for particles whose 3D orientation is constrained, such as for membrane bound particles.

Because the AP classification task is sensitive to the direction of membrane normals, it is recommended to perform few iterations of the constrained high symmetry refinement, rotational averaging and AP classification. After each iteration, classes that are not well resolved are removed and the remaining particles are again 3D refined with a high symmetry imposed to further improve the direction of their membrane normals and their location. In addition, particles that essentially merge with other particles (within 1-2 nm) as a result of the 3D refinement induced translations are removed by imposing a small exclusion distance.

#### 2.3.5 Projecting particles

Colocalization typically concerns particle sets that are located in closely apposed but separate regions, such as on pre- and postsynaptic membranes. Because the aim of the colocalization task is to determine particle distance in the lateral direction, that is along regions (such as membranes) and to minimize the contribution in the transverse direction (between regions), particles are projected onto a common 2D-like projection region. The distance between projected particles is then used for colocalization.

It is convenient to define a thin projection region that lies between particles sets whose colocalization is to be calculated. This can be accomplished by generating the 1 or few pixels thick boundary (face) of one previously selected region that directly contacts another region (Suppl. figure S7C) using basic image morphology operations. Depending on the segmented regions involved, the resulting projection region can be curved, while a previous smoothing of the segmented regions ensures that the projection region is smooth.

In cases when projection distances are small and the projection region is smooth, projections can be determined by finding the shortest distance from a point to be projected to the projection region region, as done previously (Liu et al., 2020; Chen et al., 2020). When the projection membrane is uneven, or projection distances are large, we suggest using a method whereby particles are projected along a line defined by the particle position and its membrane normal vector (Suppl. figure S7C). For particles where the line projection could not be found, the projection is determined by the shortest distance approach.

#### 2.3.6 Restricting segmented regions

Segmentation sometimes results in regions that are larger than the part occupied by particles located by the above steps. This can happen during segmentation because the extent of a particle distribution is not clear, but also deliberately, to make sure that as many particles as possible are detected. As a result, particles will appear slightly clustered within the segmented region.

As shown above, clustering minimally affects the colocalization task results when the area over which particles are distributed is covers at least 80% of the region. Furthermore, stronger clustering can be distinguished from particle interactions by the colocalization task. Nevertheless, it is advisable to remove this artificially induced clustering because it makes interpretation of colocalization results more straightforward.

For this reason, we implemented a procedure that restricts a region to the part occupied by particles. First, we generate the convex hull of a particle set, that is the smallest convex polyhedron that contains all particles. The hull is then slightly expanded to avoid that particles are located on the hull boundary. Finally, the intersection of the hull and the original region yields the restricted region. We suggest that particles comprising all retained AP classes, as opposed to one class, are used to determine complex hull because individual classes may be clustered.

## 3 Discussion

A single cellular cryo-ET dataset can provide the location of hundreds of (proteins and) protein complexes of multiple molecular types with a single nanometer precision, together with the information about their cellular environment. Mounting evidence indicates that assemblies comprising multiple species of protein complexes organized in a non-periodic manner, about one to several hundred nanometers in size, which often emerge from colocalization of nanodomains, form functional units relevant for cellular function (Biederer et al., 2017; Chen et al., 2018; Choquet, 2018; Scheefhals and MacGillavry, 2018; Zuber and Lucic, 2022). Here we present a thorough characterization of the colocalization method, a parameter-free second order bi- (multi-) variate point pattern analysis method, which analyzes and characterizes spatial relations between multiple species of protein complexes, and determines the statistical significance by a Monte Carlo method.

Numerical applications to synthetic point particle patterns designed to mimic distribution of complexes in cells showed that the method correctly detected interactions between two and between three patterns, and identified sensitivity limits. Furthermore, we provide guidelines for interpretation of results needed not only to determine whether the observed particle localization is generated by a cellular process, as opposed to a purely stochastic one, but also to determine which type of biochemical interactions and cellular processes is likely responsible for the observed particle localization.

To be able to distinguish different particle organizations relevant for biological applications, we designed several point patterns having distinct, well-defined properties that mimic different forms of particle organization that occur in biological systems. Specifically, (completely) random patterns contained particles that were randomly distributed over the entire region, such as a lipid membrane, which would be expected in systems dominated by free diffusion. Clustered patterns contained particles randomly distributed over well defined parts covering up to 80% of the entire region. These could be generated by diffusion of complexes of interest constrained by dynamic binding to scaffolding proteins loosely organized over a certain area. Another type of univariate pattern was obtained by imposing a minimum distance between otherwise randomly distributed patterns, which could arise from steric hindrance or another repulsive interaction.

Of particular importance is colocalization between two or more interacting patterns, that is particle sets whose localization depends on each other. The interaction patterns we generated fall in two categories. (i) Global interactions, where patterns are clustered over the same area, but the individual particle locations are independent. These can occur when two scaffolding processes, each binding particles of one set, are constrained to the same area. We also designed a weaker interacting variant of this pattern where the patterns partially overlap, corresponding to the situation where the two scaffolding processes are only to some extent correlated. (ii) Local interactions, where some individual particles from different sets “interact”, that is they are placed in vicinity of each other. We generate two modes, in the first mode (1-to-1) interacting particles from disjoint pairs, as expected from binary protein interactions. In the second mode (multimeric), one particle can interact with multiple particles of the other set. In both modes, only subsets of particles were involved in interactions. The other particles were placed randomly, in order to account for binding kinetics of biochemical interactions, and a possible molecular and functional heterogeneity present within particle sets.

Applications to different patterns by numerical simulations of colocalization between two patterns resulted in the correct detection of local interactions even when only 10% of particles were interacting for 3-colocalizations, 15% for 2-colocalizations in 1-to-1 and 50% in the multimeric mode. Furthermore, 3-colocalizations showed an increased sensitivity when the same subset of one particle set interacted with the other two sets.

Global interactions were detected when particles were clustered over 60% or less of the entire area, even when particle sets were partially overlapping. Furthermore, the contributions of local and global interactions were detected and distinguished from each other in cases where both interactions were present simultaneously. Regarding non-interacting sets, colocalizations where only one particle set was clustered were not significant according to the combined test, but were clearly detected by considering test results separately. Interestingly, very mild clustering (particles spread over 80% over the entire area) and small particle exclusion distance (5 nm) did not influence the results, indicating that the effects of imprecise segmentation and finite size of complexes do not affect colocalization analysis. Based on these results, we extracted guidelines for executing the colocalization task and for the interpretation of experimental results, which benefit from the possibility to change the order of particle sets and use two independent null models.

We also derived the analytical solution for colocalization of completely random and clustered patterns and showed that the small discrepancies between the analytical and numerical results can be accounted for by introducing boundary corrections. Furthermore, we modeled local interactions by introducing an interaction strength parameter. These allow a deeper understanding of the colocalization results, serve as an independent confirmation of numerical solutions and provide a quantitative characterization of interaction strength and boundary effects.

Previously, colocalization based on distances between individual particles located in nanodomains or similar well defined regions was detected by methods such as cross-correlation, nearest neighbor and Ripley’s functions in images obtained by single molecule super-resolution fluorescence microscopy, immunogold electron microscopy and cryo-ET (Tang et al., 2016; Haas et al., 2018; Rebola et al., 2019; Liu et al., 2020; Martinez-Sanchez et al., 2020; Chen et al., 2020). The design of our colocalization method reflect the imaging method properties. For example, in super-resolution fluorescence imaging, particle identity is unambiguous, but particle detection is complicated by the fluorescence background and overcounting (Haas et al., 2018; Chen et al., 2020), in contrast to cryo-ET where particles are localized with a higher precision and their cellular environment is evident, but distinguishing molecular species critically depends on the classification task. Compared to Ripley’s functions (Martinez-Sanchez et al., 2022), our method is computationally simpler, allows a straightforward extension to more than two patterns, and it is designed to detect direct or indirect biochemical interactions between individual particles. Regarding the previous implementation (Martinez-Sanchez et al., 2021, 2022), the one presented here provides much faster execution, application examples and cleaner data structures that allow immediate access to colocalized particles.

In addition, we present the colocalization workflow, which contains cryo-ET image processing tasks that follow tomographic reconstruction and precede the colocalization task. While most of these methods were used in cryo-ET before, here we specify their role within the context of the colocalization workflow. We give usage instructions, such as those for generating thin, smooth, curved regions and projecting particles onto them. Together with template-free detection and unsupervised classification, they serve to provide an optimal setup for the colocalization task.

The colocalization method allows quantitative characterization of the spatial organization of assemblies comprising multiple species of protein complexes. This organization is governed by a multitude of biochemical interactions on the background of constrained diffusion and is expected to have a functional significance. While initially developed for neuronal synapses and other cellular junctions, the colocalization method presented here can be equally applied to protein assemblies located on intracellular membranes, such as Golgi stacks, or located in other subcellular regions. Together, the colocalization allows investigations of large, non-periodic protein assemblies and is expected to bring the understanding of their function to a new level.

## 4 Materials and methods

### 4.1 Software implementation and availability

All software developed and used for this work was written in Python 3 language, it is open-source and is distributed as part of Pyto package (version 1.11, available on GitHub https://github.com/vladanl/Pyto) (Lucic et al., 2016). Pyto uses NumPy, SciPy, Jupyter Pandas scientific computing packages, as well as Matplotlib for plotting (Harris et al., 2020; Virtanen et al., 2020; Wes McKinney, 2010; Kluyver et al., 2016; Hunter, 2007).

### 4.2 Colocalization task

All point patterns (particle sets) used for the colocalization task were distributed over a rectangular region. Unless stated otherwise, this region was 400×200 pixels in size, at 1 nm pixel size. The number of particles was 300, 250 and 50, for sets *P, Q* and *R*, respectively. All these were in the range of the particle numbers and surface areas encountered in the previously analyzed experimental data (Martinez-Sanchez et al., 2021). The following point patterns were generated:

- Random: Completely random over the entire area
- Clustered: Random over a region covering 50%-80% of the entire area
- Randomly distributed with a minimum exclusion distance between points
- Globally interacting patterns: Pairs of globally interacting patterns consisted of two clustered patters that completely, or partially overlapped. The partial overlap area was 705-80% of the cluster area.
- Locally interacting patterns, 1-to-1 mode (Suppl. figure S1B left): Pairs of locally interacting patterns, consisting of a fixed and the dependent patterns, were generated in the following way. A completely random or clustered pattern was used as the fixed pattern and a randomly chosen subset of the specified size was chosen as the interacting subset of the fixed pattern. Then, one point of the dependent set was placed within the colocalization distance to each of the interacting subset point, resulting in disjoint pairs of interacting points. The remaining dependent pattern points were distributed randomly. The size of the interacting pattern was 10% - 30% of the fixed pattern size and the colocalization distance ranged from 5 to 15 nm.
- Locally interacting patterns, multiple mode (Suppl. figure S1B right): Pairs of locally interacting patterns, consisting of a fixed and the dependent patterns, were generated in the following way. A completely random or clustered pattern was used as the fixed pattern and a randomly chosen subset of the specified size was chosen as the interacting subset of the fixed pattern. Then, each of a specified number of the dependent set points was randomly assigned to any of the interacting subset points, in a completely independent manner, and placed within the colocalization distance to the selected interaction subset point. This generally results is some of the interacting subset points having no interacting partners (from the dependent set), while other interacting ponts having one or more interacting partners. Fraction of interacting points from the fixed and the dependent patterns was independent from each other and was between 30% and 70% of the total number of points. The colocalization distance ranged from 5 to 15 nm.
- Interacting patterns for 3-colocalizations, common interacting subset variant (Suppl. figure S1C left): The three interacting subsets consist of two pairs of locally interacting patterns (mode 1-to-1) with a common fixed pattern, together making one fixed and two dependent patterns. In this variant, the two interacting patterns had the same interacting subset of the fixed pattern.
- Interacting patterns for 3-colocalizations, independent interacting subset variant (Suppl. figure S1C right): The three interacting subsets consist of two pairs of locally interacting patterns (mode 1-to-1) with a common fixed pattern, together making one fixed and two dependent patterns. In this variant, the interacting subsets of the fixed pattern, used for the two interacting patterns, were independent of each other.

Random generator seeds were different for all random patterns that were generated, to ensure that all random patters were independent from each other. To ensure the reprehensibility of the results, all seeds were fixed, they started from an arbitrary one (123456) and generally increased by 1.

Here we reimplemented in Pyto package the colocalization calculation code, which was previously in PySeg and Pyto packages, and added the code that generates particle patterns described above. Calculations of distances between particles, the core of the colocalization calculations, now uses Scipy functions for Euclidean distance calculation (Virtanen et al., 2020). All operations are in real (pixel space), as opposed to polygons used in computer vision, which avoids conversions to and from (pixel based) MRC image format, Classes that store and handle all particles and tomogram info, such as particle ids, coordinates and class labels, as well as colocalization results now use Pandas tables (Wes McKinney, 2010). Furthermore, calculating a 3-colocalization together with the related 2-colocalization is now more efficient because the random particle patterns generated for Monte Carlo simulations are reused. All this contributed to more than 10 times faster execution, and the cleaner data structures allow immediate access to the results including the specification of particles that actually colocalize.

An example that uses Jupyter notebooks (Kluyver et al., 2016), showing point pattern generation, numerical calculations of colocalization and the results analysis for all graphs presented here is provided in the Pyto package (see Software availability, below).

### 4.3 Analytic solution and heuristic parameters

The boundary correction parameters, *ξ* and *α*, were used only in cases for which they were designed. That is, for colocalization *P* -*Q, ξ <* 2 was used when boundaries of *P* and *Q* areas coincided and *α >* 0 when the area of *P* was inside the area of *Q* (Suppl. figure S1). The value *ξ* = 1.96 was used on all graphs unless stated otherwise. It was determined by fitting the numerical results for the colocalization between two completely random patterns (Figure 2A) and it was used for all other cases when the area boundaries of *P* and *Q* were the same, except for the extremely elongated area (Suppl. figure S2A).

The interaction parameter *β* was used to fit the numerical results for locally interacting patterns. Values

*β >* 1 indicate attraction between points.

### 4.4 Implementation of other tasks

Smoothing segmented regions by morphological functions and restricting segmented tasks by convex hull were implemented in Pyto package using functions from Scipy (Virtanen et al., 2020).

Particle projections on a region based on the minimal distance to the region and along the membrane normal line at the particle positions are implemented in Pyto package. The latter is particularly useful for membrane bound complexes where the projection line should be perpendicular to the membrane.

Projection lines are defined by the location of the particle and the line direction, where the latter is determined by the previously calculated Euler angles of the particle. For example, if Euler angles were obtained by Relion (Bharat and Scheres, 2016) 3D refinement with a high C symmetry, the projection line direction is determined by Tilt and Psi angles. More generally, the last two of the active, intrinsic, zyz Euler angles convention are required. The projection point is determined as the closest grid point of the region that is within a specified distance to the projection line. This distance is determined for each tomogram, as the one for which the highest number of distinct projection points are found.

Examples of smoothing regions by morphological operations, restricting regions by convex hull and particle projection, accessible via Jupyter notebooks (Kluyver et al., 2016), are provided in the Pyto package pyto/examples (see Software availability, below).

Morse theory based particle detection and particle classification by Affinity propagation were used as previously implemented in the PySeg package (Martinez-Sanchez et al., 2020).

## Appendix

### Derivation of analytical solutions

#### Random patterns

We now derive the analytical solution for the expected number of colocalizations between particle sets *P* and *Q*, comprising *N*_*P*_ and *N*_*Q*_ particles respectively, randomly distributed on flat surfaces of area *A*_*P*_ and *A*_*Q*_ with overlap area *A*_0_. The probability that a given particle from *Q* is not located in the neighborhood *a*(*d*) of a particle from *P* is: 1 *− a*(*d*)*/A*_*Q*_, where *a*(*d*) is the area of the neighborhood parametrized by distance *d*. Because particles of *Q* are independent (their position is randomly distributed) the probability that none of the particles from *Q* is in the neighborhood of a given particle from *P* is:

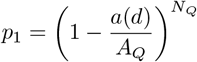

The probability that one or more particles of *Q* are in the neighborhood of a given particle from *P* is simply 1 *− p*_1_. This is by definition the probability that a given particle from *P* is colocalized with set *Q*.

Multiplying this probability with the expected number of particles of *P* that contribute to the colocalization (those located in the overlap of areas where *P* and *Q* are distributed) yields the expected number of particles of *P* that are colocalized with *Q*, which is by definition the number of colocalizations (*E*_*PQ*_):

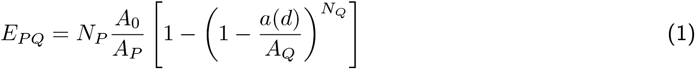

We note the following properties of this equation. First, the above derivation requires that *P* and *Q* are independent and that particles of *Q* are randomly distributed within area *A*_*Q*_. However, it places no constraints on the distribution of *P*. In other words, the equation is valid for any (e.g. random or clustered) distribution of *P*, provided that the above two constraints are satisfied. Second, although we are here concerned with particles distributed on 2D surfaces, the equation is valid for any number of spatial dimensions, provided that *a*(*d*) is adequately defined.

More generally, because colocalization of *N* particle sets (*P, Q*_1_, *Q*_2_, … *Q*_*N−*1_) requires that at least one particle from each of the *Q*_*i*_ sets is localized in the neighborhood of a particle of *P*, the expected number of colocalizations can be written as

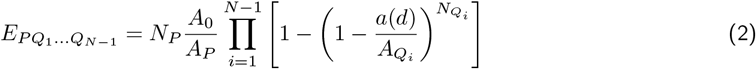

In the limit when *A*_*Q*_ ≫ *a*(*d*) and for arbitrary *N*_*Q*_ *>* 1, Eq. 1 can be converted to a simpler form:

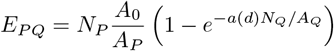

#### Boundary corrections and interaction

To obtain a solution applicable to a wider set of problems, we first investigate the effect of the boundaries of *A*_*P*_ and *A*_*Q*_. Eq. 1 is valid in the limit where the areas over which particles are distributed approach infinity and the number of particles per unit area is finite. For particles distributed on 2D surfaces and ignoring the influence of the boundaries of areas *A*_*P*_ and *A*_*Q*_, the neighborhood is a circle of radius *d*, so the neighborhood area is *a*(*d*) = *πd*^2^.

However, when the areas of *P* and *Q* are finite and coincide with each other, the neighborhood of particles from *P* that are located at a distance smaller than *d* to the area boundary is the intersection of the circle of radius *d* centered at the particle location and the area of *Q*, effectively reducing the neighborhood area *a*(*d*) (Suppl. figure S1A). This effect becomes more prominent as the ratio of the boundary length and the area of *Q* increases. In the extreme case, when the area of *Q* goes to to 0, the colocalization becomes 1-dimensional.

The following form for the neighborhood area *a*(*d*) is correct for infinite areas of *P* and *Q* in both 1 and 2 dimensions, depending on a heuristic parameter *ξ*:

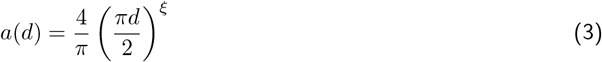

as *a*(*d*) = 2*d* for *ξ* = 1 and *a*(*d*) = *πd*^2^ for *ξ* = 2. Because *a*(*d*) monotonically increases with *ξ* (termed effective dimensionality), we can account for the presence of the boundary by setting *ξ <* 2. Therefore, substituting Eq 3 in Eq 1 provides an approximate solution for the number of colocalizations *E*_*PQ*_.

When the area of *Q* is completely inside and surrounded by the area of *P*, the Eq. 1 may underestimate the number of colocalizations, because of another type of boundary effect. Namely, particles of *P* located outside the area of *Q* at a distance smaller than *d* to the boundary may also contribute to the number of colocalizations (Suppl. figure S1A), but they are not accounted for in Eq. 1 by the term that specifies the expected number of particles of *P* contributing to the colocalization (*N*_*P*_ *A*_0_*/A*_*P*_). Because this effect increases with *d*, we account for it by introducing a multiplicative factor 1 + *αd* that depends on parameter *α*, where *α >* 0 effectively increases the number of particles of *P* located in the overlap area.

To model interaction between individual points, we introduce interaction parameter *β*, where *β >* 1 effectively increases the neighborhood area *a*(*d*) and thus number of colocalizations *E*_*PQ*_.

Taking all these corrections together, the number of colocalizations that approximates the influence of boundaries to the first order, in 2D, becomes:

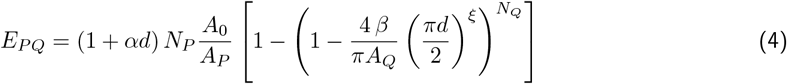

where parameters *ξ* and *α* depend on the shape of the area where particles are distributed, and *β* signifies the interaction strength. These parameters do not depend on the colocalization distance *d* and they need to be determined heuristically.

## CRediT authorship contribution statement

D.H.O.-B.: Methodology; Software; Investigation; Writing - Review & Editing. A.M.-S.: Methodology; Writing - Review & Editing; Supervision; Funding acquisition. V.L.: Conceptualization; Methodology; Software; Formal analysis; Investigation; Writing - Original Draft; Writing - Review & Editing; Supervision; Funding acquisition.

## Generative AI statement

During the preparation of this manuscript generative AI and AI-assisted technologies were not used at all.

## Software and data availability

All software developed and used for this work is open-source and distributed as a part of Pyto package (version 1.11), available on GitHub https://github.com/vladanl/Pyto) (Lucic et al., 2016).

All colocalization task data and code used here are provided as the colocalization task example in the Pyto package. It shows point pattern generation and numerical calculations in notebook pyto/examples/colocalization/colocalization task/make patterns.ipynb, and the colocalization results analysis for all graphs presented here in notebook pyto/examples/colocalization/colocalization task/analysis.ipynb.

Usage examples for other colocalization workflow tasks introduced here are also distributed as a part of Pyto package (in pyto/examples/colocalization/other colocalization workflow tasks/). These include Smoothing by morphological operations (morphological smoothing.ipynb), Particle projection (particle projection.ipynb) and Restricting regions by convex hull (convex hull.ipynb).

All examples are presented in Jupyter notebooks (Kluyver et al., 2016), they are self contained and can also be used to process external data.

## Acknowledgments

We would like to thank Wolfgang Baumeister and Juergen Plitzko for their support, and Gabriela J. Greif for critical reading of the manuscript.

## Supplementary information

The following supplementary Information is available for this paper: Suppl. figures S1–S7

## Supplementary Material

This PDF includes Suppl. figures S1–S7

**Figure S1.**
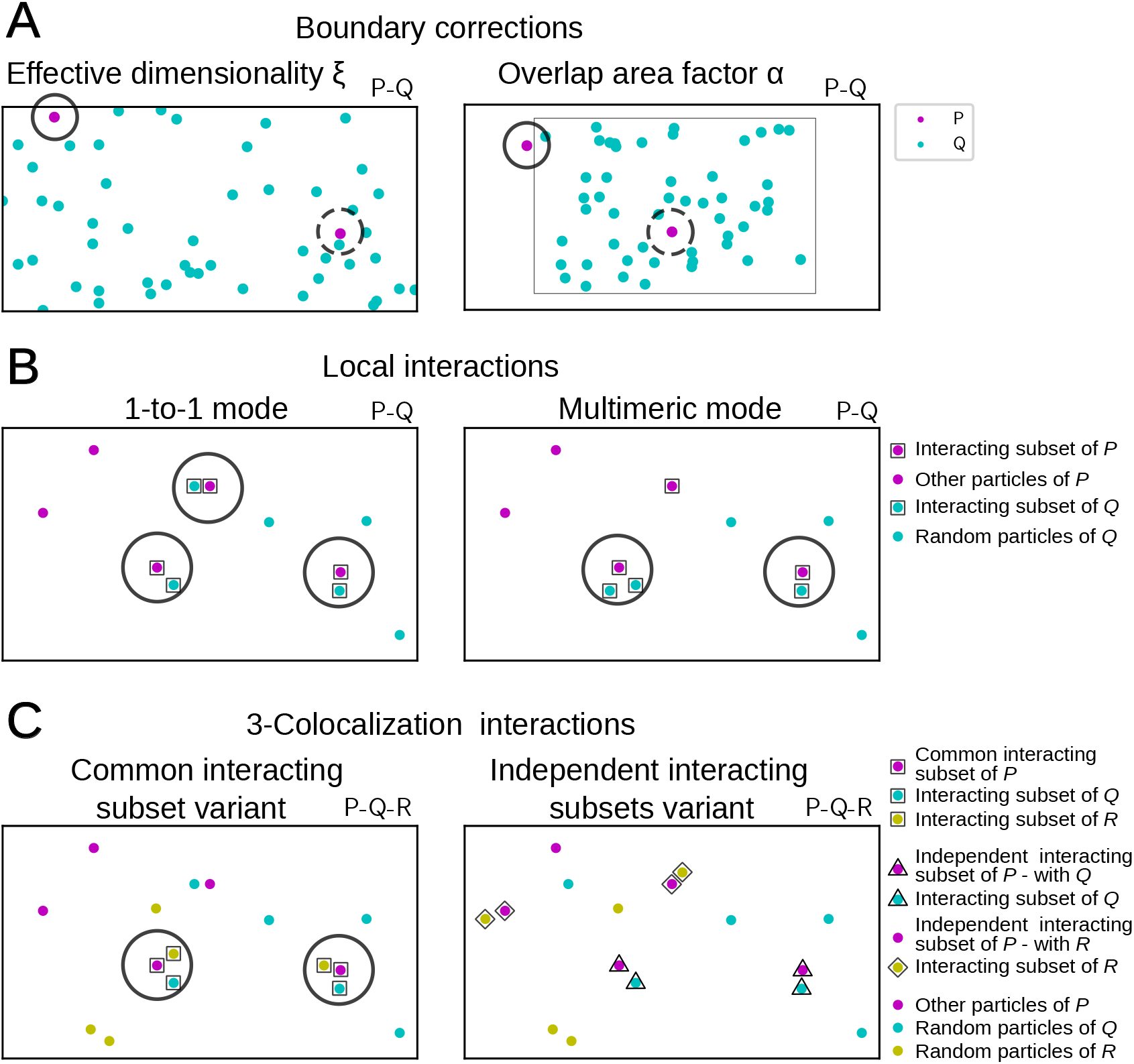
Boundary corrections and colocalization modes. (A) Boundary corrections. On the right panel, solid circle shows the neighborhood of a particle of set *P* that contains a region outside the image, where no particle of *Q* can be found. This causes the overestimate by the analytical formula that is corrected by effective dimensionality parameter *ξ*. On the left panel, solid circle shows the neighborhood of a particle of set *P* located outside the intersection of areas of *P* and *Q*. This causes an underestimate by the analytical formula that is corrected by the area overlap parameter *α*. On both panels, dashed circles show neighborhoods that are correctly taken into account in the analytical formula. (B) Interaction patterns consisting of a fixed pattern *P* and dependent pattern *Q* generated in 1-to-1 (left) and multimeric mode (right). Particles belonging to the interacting subsets are enclosed in squares. Circles show colocalized particles. (C) 3-colocalization interaction patterns. The left panel shows the common interacting subset variant, where particles belonging to the interacting subset of *P* and their interacting partners from sets *Q* and *R* are enclosed in squares. The right panel shows the independent interacting subsets variant, where particles of the first and the second interacting subset of *P* and their interacting partners from *Q* and *R* are enclosed in triangles and diamonds, respectvely. Circles show colocalized particles for *P* -*Q* -*R* colocalization.

**Figure S2.**
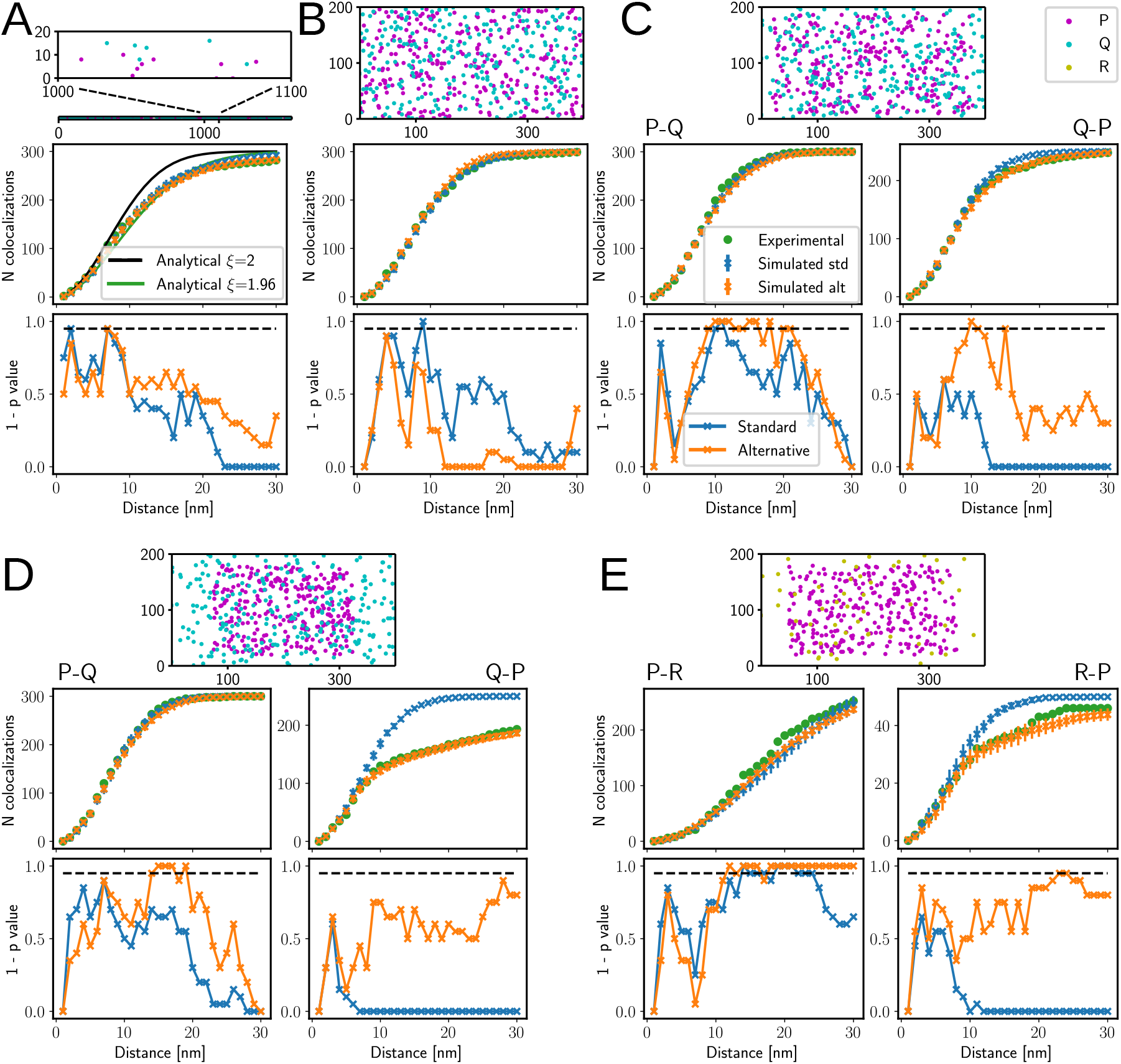
Colocalizations between non-interacting sets. (A) Completely random sets *P* and *Q* located over an extended area, (B) Completely random set *P* and *Q* with 5 nm particle exclusion distance. (C) Colocalization between *P* clustered over 80% over the entire area and *Q*. (D) Colocalization between *P* clustered over 50% over the entire area and *Q*. (E) Colocalization between *P* clustered over 60% over the entire area and *R*. (A-E) Particle distribution is shown on top, number of colocalizations in the middle and the p-values at the bottom. Horizontal dashed line shows 0.05 confidence limit. (C-E) The colocalization order is specified above graphs. Legends shown on C apply to all panels.

**Figure S3.**
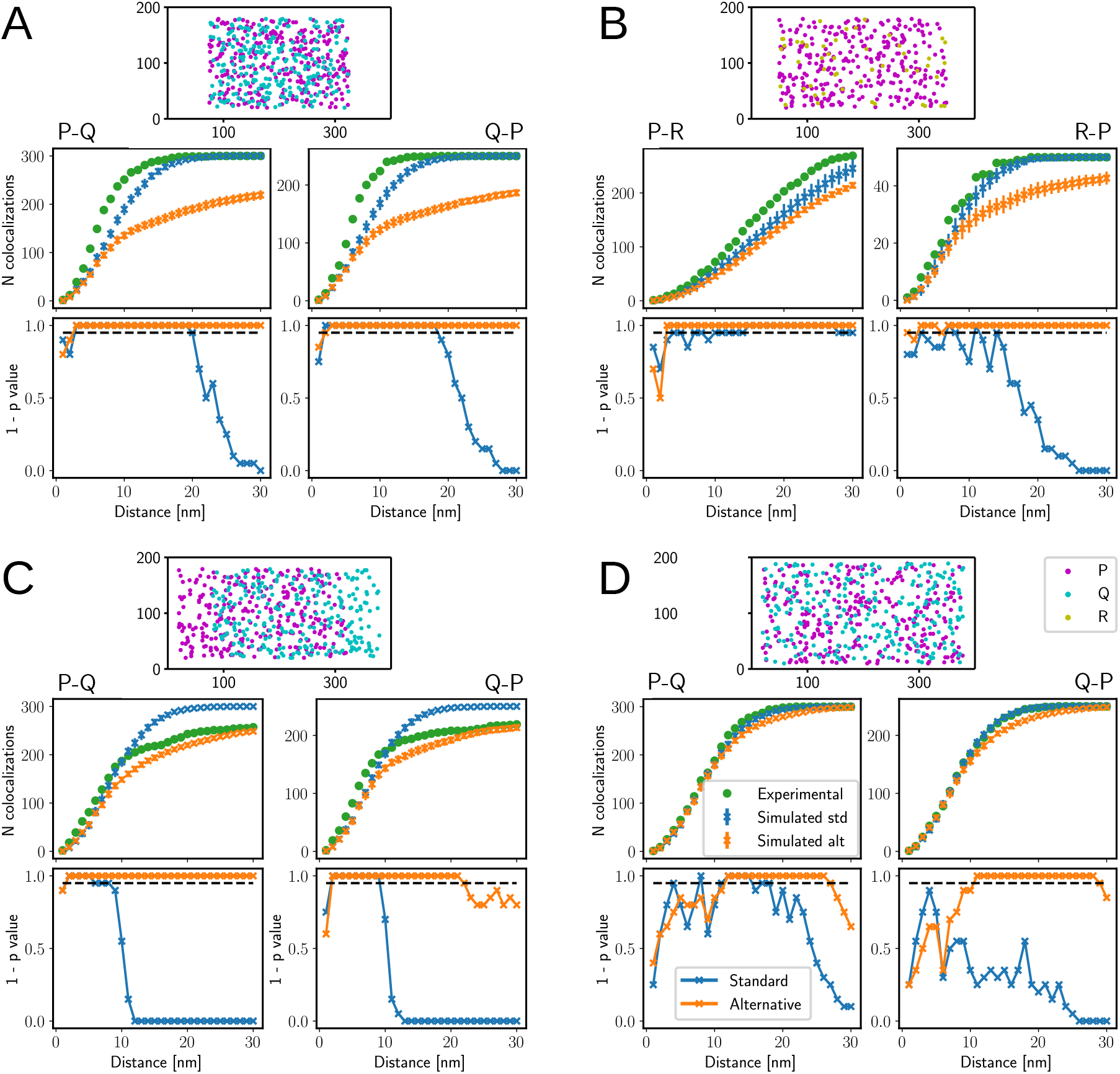
Colocalizations between globally interacting particle sets. (A) Sets *P* and *Q*, clustered on completely overlapping regions covering 50% of the entire area. (B) Sets *P* and *R*, clustered on completely overlapping regions covering 60% of the entire area. (C) Sets *P* and *Q*, clustered on partially overlapping regions, both covering 60% of the entire area. (D) Sets *P* and *Q*, clustered on completely overlapping regions covering 80% of the entire area. (A-D) Particle distribution is shown on top, number of colocalizations in the middle and the p-values at the bottom. Horizontal dashed line shows 0.05 confidence limit. The colocalization order is specified above graphs. Legends shown on D apply to all panels.

**Figure S4.**
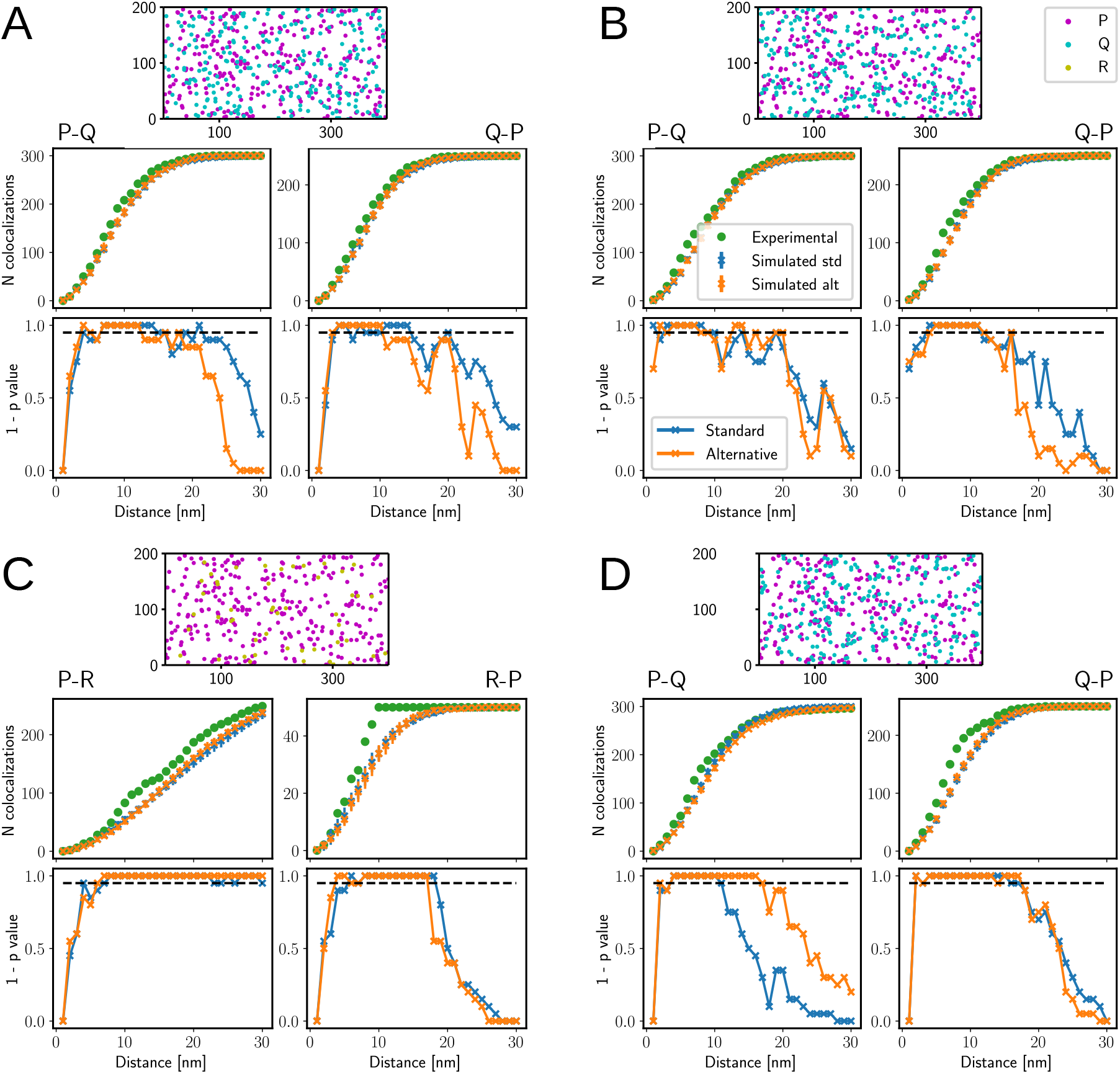
Colocalizations between locally interacting particle sets. (A) Sets *P* and *Q*, 1-to-1 mode, interacting subset contained 15% of *P*, interaction distance 10 nm. (B) Sets *P* and *Q*, 1-to-1 mode, interacting subset contained 15% of *P*, interaction distance 5 nm. (C) Sets *P* and *R*, 1-to-1 mode, interacting subset contained 15% of *P*, interaction distance 10 nm. (D) Sets *P* and *Q*, multimeric mode, interacting subset contained 50% of *P* and *Q*, interaction distance 10 nm. (A-D) Particle distribution is shown on top, number of colocalizations in the middle and the p-values at the bottom. Horizontal dashed line shows 0.05 confidence limit. The colocalization order is specified above graphs. Legends shown on B apply to all panels.

**Figure S5.**
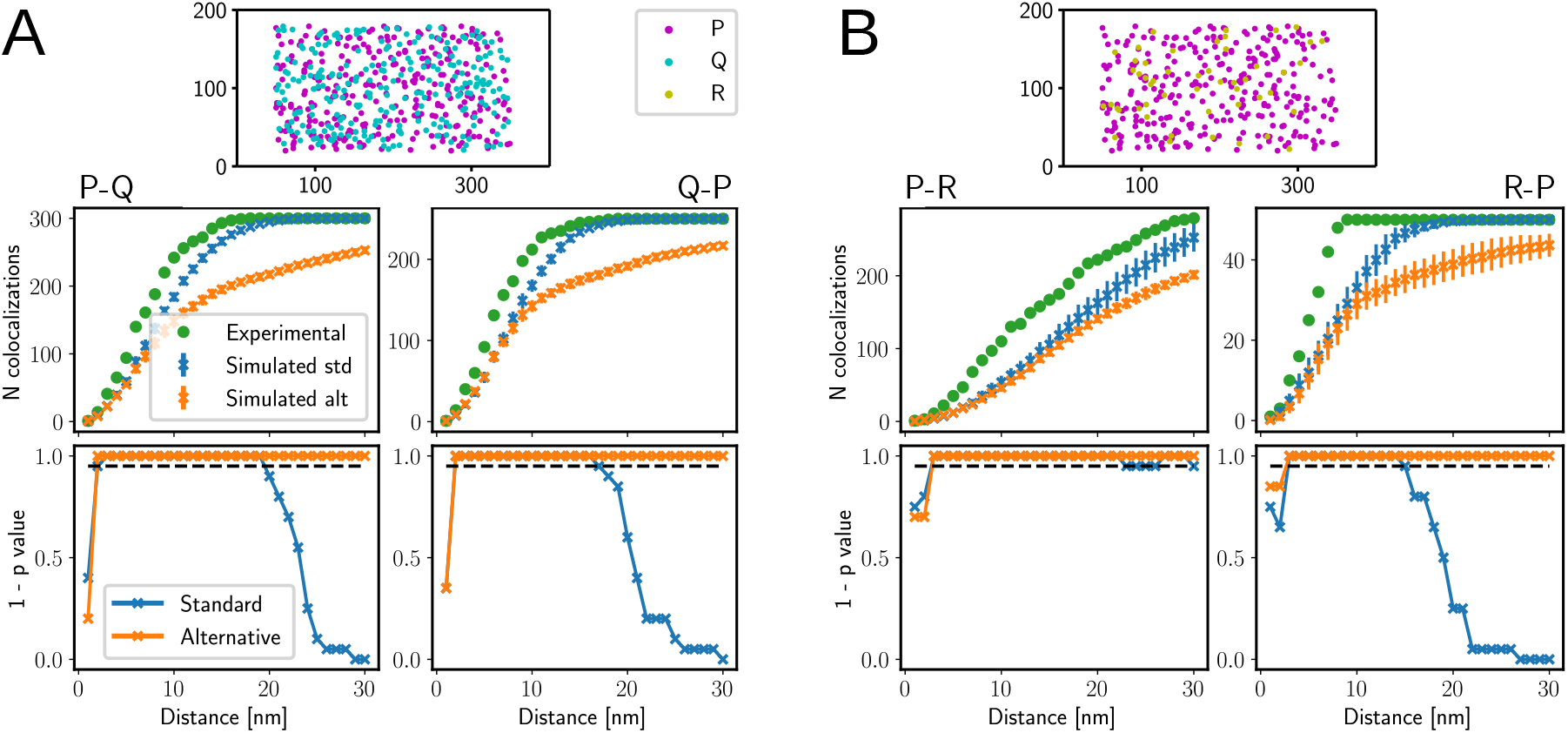
Colocalizations between particle sets that interact globally and locally. (A) Sets *P* and *Q*, clustered over 60% over the entire area, 1-to-1 mode, interacting subset contained 15% of *P*, interaction distance 10 nm. (B) Sets *P* and *R*, clustered over 60% over the entire area, 1-to-1 mode, interacting subset contained 15% of *P*, interaction distance 10 nm. (A-B) Particle distribution is shown on top, number of colocalizations in the middle and the p-values at the bottom. Horizontal dashed line shows 0.05 confidence limit. The colocalization order is specified above graphs. Legends shown on A apply to both panels.

**Figure S6.**
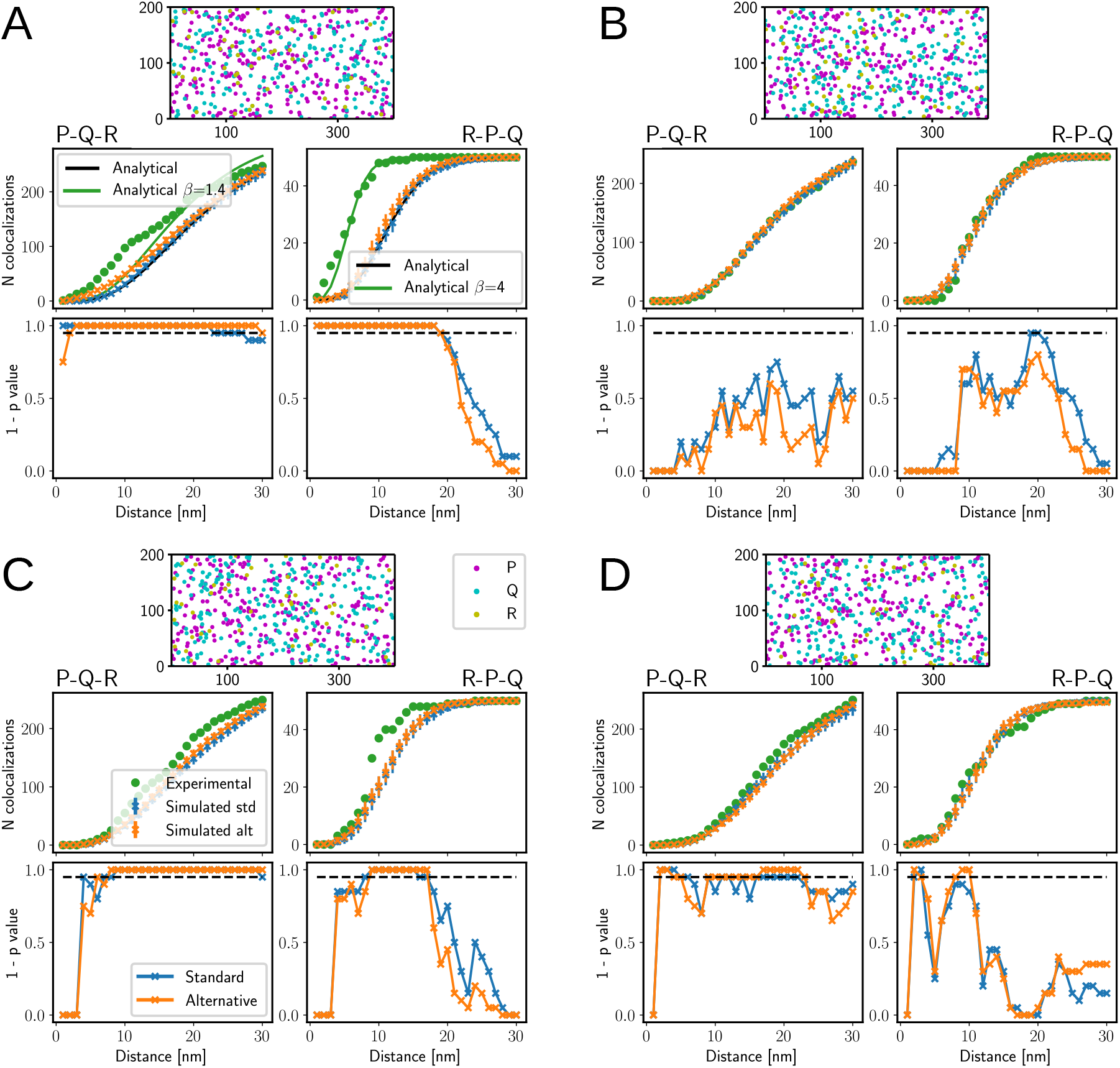
3-Colocalizations. (A) Local interaction, common interacting subset variant, the subset contains 15% particles of *P*. (B) Completely random, independent sets. (C) Local interaction, independent interacting subset variant, the subset containes 15% of particles of *P*. (D) Local interaction, independent interacting subset variant, the subset contains 10% of particles of *P*. (A-D) Particle distributions are shown on top, number of colocalizations in the middle and the p-values at the bottom. In all interacting cases, 1-to-1 mode was used and the interaction distance was 10 nm. Horizontal dashed line shows 0.05 confidence limit. The colocalization order is specified above graphs. Legends shown on C apply to all panels.

**Figure S7.**
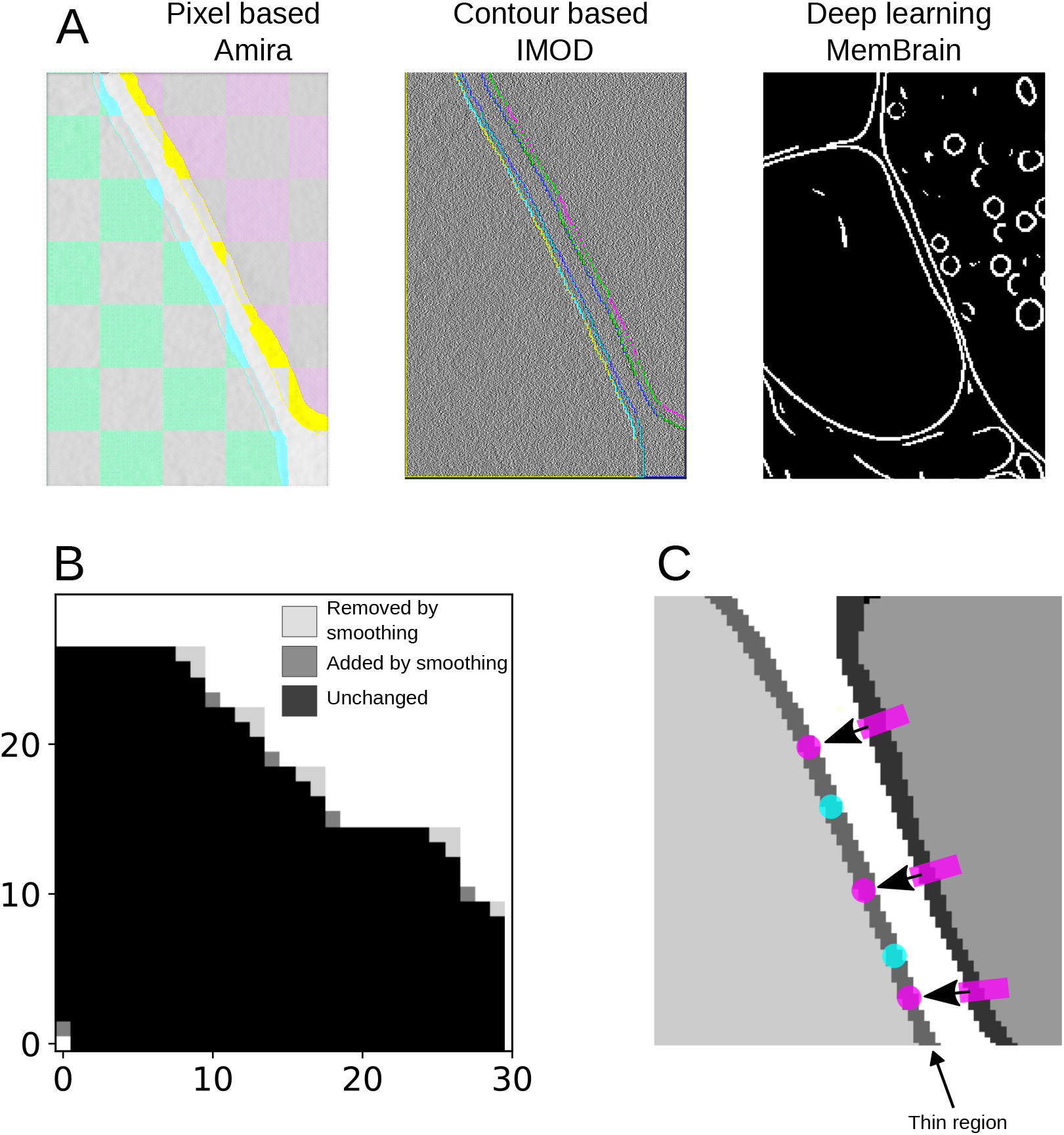
Other colocalization workflow tasks. (A) Pixel-based segmentation by Amira™(left), contour-based segmentation by MOD (center) and deep learning segmentation (right) (Stalling et al., 2005; Kremer et al., 1996; Lamm et al., 2022). (B) Smoothing a segmentation obtained at a higher binning by morphological operations. (C) Projecting particles on a thin region. Rectangles show the initial and circles (of the same color) the final particle positions. Arrows associated with particles show the projection direction.

